# Mechanisms of macular oedema development and therapeutic response: An in-silico modelling study

**DOI:** 10.64898/2026.02.17.706117

**Authors:** Pradeep Keshavanarayana, Emmeline Brown, Philip Luthert, Rebecca Shipley, Simon Walker-Samuel

## Abstract

**Purpose:** To investigate the interplay between biomechanics, fluid dynamics, and solute transport in Diabetic Macular Oedema (DMO) using a mechanics-based computational model, aiming to elucidate mechanisms behind variable treatment outcomes.

**Methods:** We developed a multiphysics model of the retina within a porous media framework. The model integrates OCT-derived geometry, vascular leakage, retinal biomechanics (including Müller cell fibre architecture), retinal pigment epithelium (RPE) function, and anti-VEGF transport. We simulated oedema development and therapeutic response by varying these parameters systematically.

**Results:** Model results showed that active RPE pumping is essential for maintaining retinal dehydration. Our simulations revealed a critical trade-off related to Müller cell architecture: the physiological ‘z-shaped’ orientation protects against oedema but impedes anti-VEGF drug delivery to leaky vessels. In contrast, a pathological, vertical Müller cell alignment increases oedema susceptibility but allows for a faster therapeutic response due to improved drug diffusion.

**Conclusions:** Müller cell orientation presents a trade-off between biomechanical protection and therapeutic efficacy, offering a novel mechanistic explanation for the variable patient responses to anti-VEGF therapy observed clinically. This in-silico framework is a powerful tool for dissecting DMO pathophysiology and has the potential to guide the development of personalised treatment strategies.

## 1 Introduction

Diabetic macular oedema (DMO) is one of the leading causes of vision loss and a growing global health concern, affecting around 20 million people worldwide [1]. Prolonged high blood glucose levels (hyperglycaemia) damage retinal vasculature, leading to diabetic retinopathy, projected to affect 45 million individuals by 2045 [2], and frequently progresses to DMO. The prevalence of DMO varies widely, ranging from 4.2% to 14.3% in people with type 1 diabetes mellitus (T1DM) and 1.4% to 5.57% in people with type 2 diabetes mellitus (T2DM) [3].

DMO is characterised by abnormal accumulation of fluid in the macula, the central retinal region responsible for high acuity vision. Retinal homeostasis relies on a delicate balance between osmotic and hydrostatic pressures, modulated by the blood-retinal barrier (BRB). The BRB is composed of an inner barrier (formed by tight junctions in the retinal blood vessels) and an outer barrier (formed by the retinal pigment epithelium, or RPE). The BRB’s primary role is to strictly control the passage of water, ions, and proteins. In diabetic patients, sustained high blood sugar (hyperglycaemia) damages this barrier. The compromised BRB leads to leaky retinal vessels and impairs the RPE’s ability to pump fluid away. As a result, fluid and solutes build up in the macula, causing it to swell and thicken—the hallmark of DMO. This process is often accompanied by structural changes, including retinal stiffening. For example, the breakdown of the BRB can trigger Müller cells to deposit excess extracellular matrix (ECM) proteins, while hyperglycaemia promotes the formation of advanced glycation end products (AGEs) [4, 5].

Additionally, the cellular architecture of the retinal cells, particularly the Müller glial cells, significantly influences the retinal biomechanical behaviour. These cells span the entire retinal thickness, acting as structural fibres that provide mechanical support and guide fluid transport [6, 7]. In a healthy macula, Müller cells exhibit a characteristic ‘z-shaped’ orientation. However, in pathological states like DMO, these cells can undergo reactive changes, altering their morphology to a more vertical alignment [7]. A combination of these changes leads to oedema development, increasing the thickness of the retina around the macula, and reducing visual acuity.

Current treatments, including anti-VEGF injections, target vascular leakage but show inconsistent efficacy across patients and often require repeated, costly interventions. Anti-VEGF agents function by either binding to vascular endothelial growth factor (VEGF) or its receptor, thereby preventing it from activating VEGF on the surface of endothelial cells [3]. This inhibition reduces neovascularisation and vascular permeability, which are key drivers of fluid accumulation in DMO. Several anti-VEGF drugs are currently available for clinical use, including bevacizumab (Avastin), ranibizumab (Lucentis), aflibercept (Eylea), and faricimab (Vabysmo), which vary in their molecular weight, and binding affinity. Patients exhibit variable responses to these treatments, with some showing significant improvement in retinal thickness and visual acuity, while others experience minimal benefits [3]. Notably, DMO progression and therapeutic outcomes are found to be influenced by patient-specific factors such as retinal geometry and the location of leakage in the vascular tree [8].

Current clinical practice primarily classifies DMO by measuring central macular thickness (CMT) or central subfield thickness (CST) using OCT imaging. However, this approach offers limited insight into the underlying physical and transport phenomena and is prone to inter-observer variability. To address these limitations, recent advancements in artificial intelligence (AI) have introduced models capable of automatically detecting, segmenting, and quantifying disease-related biomarkers from retinal images with expert-level accuracy [9, 10, 11]. By providing rapid, objective, and reproducible analysis, these AI tools help clinicians make more informed and timely therapeutic decisions. We have developed a physics-informed generative adversarial network (PI-GAN) based computational framework, incorporating physical principles of vascular network topology and fluid dynamics, for generating anatomically realistic synthetic OCT images of the retina [12]. While this model currently reproduces detailed vascular architecture and simulates diabetes-induced phenomena such as vessel occlusion and resulting redistribution of flow, it is under development for clinical translation. However, such AI models are data-dependent and require extensive longitudinal data, which are difficult to obtain, given the ethical constraints surrounding ophthalmological data.

Multiphysics modelling of tissue biomechanics is an established field with an abundance of modelling frameworks. However, its application in ophthalmology has been relatively limited, and broader clinical translation is hampered by challenges in parameterising, testing, and validating models against clinical data [13]. Several attempts have been made to develop physics-based models of the retinal physiology and pathology [14, 15]. While they offer causal insights, they have not been able to accurately predict disease progression due to limited understanding of all the biomechanical mechanisms at play, lack of parameter values governing retinal mechanochemical behaviour, and their failure to incorporate anatomically accurate geometry and integrate complex multiphysics phenomena. The recent work by Ruffini et al. [6] presents a novel multiphasic approach to model DMO, including retinal biomechanics, fluid dynamics and solute transport. However, this model does not incorporate anatomically accurate retinal geometry or vascular leakage, hindering its applicability to patient-specific DMO progression and clinical translation. Additionally, several research groups have developed mathematical models of solute transport in the eye, including anti-VEGF drugs, to improve the efficacy of ocular therapies [15, 13, 16, 14]. Despite these advances, state-of-the-art mathematical models typically focus on isolated processes occurring in the retina, withholding the complete picture of how vascular, biomechanical, and transport mechanisms interact, causing DMO.

We hypothesise that patient-specific retinal vasculature distribution and biomechanics drive variation in oedema progression. This variation, challenging to interpret clinically, could be analysed through multiphysics modelling to enable a better understanding of DMO and guide personalised treatment strategies. To test the hypothesis, in this article, we present a multiphysics computational model of the retina that integrates OCT-inspired geometry, vascular leakage, retinal biomechanics, fibre distribution morphology, fluid dynamics, and solute transport to study oedema development and anti-VEGF therapy response.

Overall, the primary innovation of this study lies in the development of a biomechanical modelling framework for the retina that integrates the complex interactions between vascular leakage, RPE function, retinal biomechanics, anatomically realistic geometry, and solute transport. This model offers novel insights into the mechanisms underlying oedema formation and therapeutic response which are currently inaccessible via experimental or clinical imaging methods alone. Specifically, our model predicts a critical trade-off whereby the physiological architecture of Müller cells, while providing biomechanical protection against oedema, simultaneously hinders the delivery of therapeutic agents, offering a new perspective on variable treatment responses. While the current framework is focused on elucidating general mechanistic trends rather than providing patient-specific predictions at this stage, it establishes a foundation that could be extended for such purposes in the future, paving the way for personalised treatment strategies.

## 2 Methods

### 2.1 Mathematical model

We have developed a mathematical model of the retina, treating it as a multiphasic deformable porous medium. This framework models the tissue as a mixture of three primary constituents: a solid matrix, an interstitial fluid (solvent), and dissolved solutes. The fundamental physical assumptions of the model are based on the principles of mixture theory. Each constituent is treated as a continuous, co-occupying medium governed by its conservation laws for mass and momentum. We assume that there is no generation or loss of mass for the solid phase; that is, we do not account for cell proliferation, death, or extracellular matrix remodelling. Interactions between the constituents are purely mechanical and chemical: momentum is exchanged via pressure and frictional drag forces, and solutes can undergo chemical reactions, such as the binding of anti-VEGF to VEGF. The multiphasic approach is necessary because the retina is an intricate, hydrated tissue where mechanical forces, fluid flow, and solute transport are tightly coupled. Modelling the tissue as *deformable* allows us to simulate the characteristic swelling and changes in retinal thickness that define oedema, which arise from the accumulation of fluid within the porous solid matrix. Biologically, the solid phase represents the retinal tissue, comprising an extracellular matrix and embedded fibres that capture the structural role of Müller cells; the fluid phase represents the interstitial fluid; and the solutes represent molecules such as albumin and anti-VEGF drugs. The model incorporates the interactions between these components, following the multiphasic material theory [17], implemented in FEBio (Version 4.10), an open source finite element solver [18]. While we reproduce some of the governing equations here for completeness, we refer the reader to prior works for detailed derivations [17, 18].

#### 2.1.1 Governing equations

The total volume of the mixture is occupied by the solid matrix, the interstitial fluid (solvent), and the dissolved solutes. Therefore, the volume fractions (the ratio of each constituent’s volume to the total volume) must sum to one:

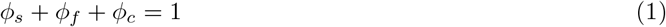

where *ϕ*_*s*_ is the solid volume fraction, *ϕ*_*f*_ is the fluid volume fraction, and *ϕ*_*c*_ is the solute volume fraction. However, for solutes like albumin and anti-VEGF drugs dissolved in the interstitial fluid, the volume they physically occupy is negligible compared to the solvent. Therefore, a standard simplification in mixture theory is to assume that the fluid and solid phases fill the entire volume:

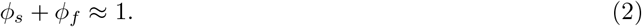

While the solute’s volume is considered negligible, its concentration and resulting osmotic pressure are fundamentally important to the model. The system is represented by three unknowns: displacement **u**, fluid pressure *p* and solute concentration per unit volume of the solution *c*. However, to solve the governing equations following the finite element method, we modify the variables as follows: The solute concentration is modified as

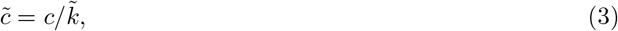

where 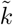 is the effective solubility coefficient and 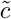 is the effective solute concentration. Here, 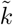 represents the reduction in solute concentration due to pore size and charge effects. The fluid pressure is modified as

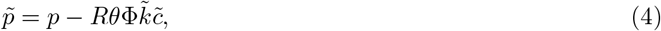

where 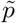 is the effective fluid pressure, *R* is the universal gas constant, *θ* is the absolute temperature and Φ is the osmotic coefficient. 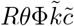 represents the osmotic pressure due to the solute. The fluid pressure is modified to account for the osmotic pressure caused by the solute. Thus, the unknowns are now displacement **u**, effective fluid pressure 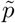 and effective solute concentration 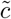.

Further, the total stress in the mixture is given by the Cauchy stress tensor

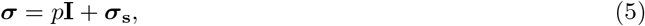

where *p* is the fluid pressure, ***σ***_*s*_ is the solid stress and **I** is the identity tensor. Writing the balance of linear momentum for the mixture, we have,

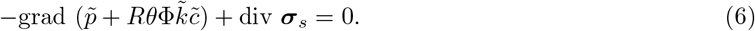

Further, the conservation of mass for the mixture is given by

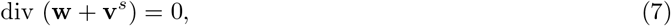

where **w** is the fluid flux relative to the solid and **v**^*s*^ is the solid velocity. Similarly, the conservation of mass for the solute, including mass generation due to chemical reactions, is given by

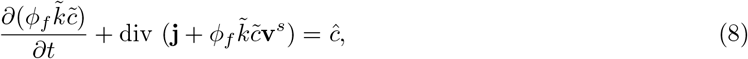

where **j** is the solute flux relative to the solid, 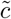 is the effective solute concentration, and *ĉ* is the rate of mass generation of the solute per unit volume. The first term on the left-hand side represents the rate of change of solute concentration over time, making this a transient equation. Writing the linear momentum balance for fluid and solute phases, we can obtain the relation between 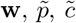 and **j** as follows:

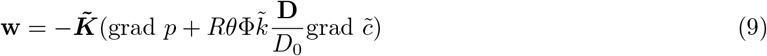

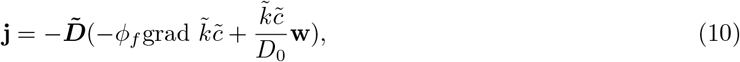

where 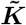 is the hydraulic permeability tensor, 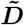 is the diffusion tensor and *D*_0_ is the reference diffusivity. Note that Equation 9 and Equation 10 reduce to Darcy Law and Fick’s law for pressure driven and solute concentration driven flows respectively. Readers are referred to the FEBio theory manual for detailed derivation and discussion of the above equations [18].

Finally, we assume that the solid stress is given by

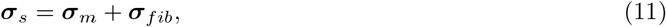

where ***σ***_*m*_ is the matrix stress and ***σ***_*fib*_ is the fibre stress. In the current article, we assume an isotropic nearly incompressible Neo-Hookean material model for the matrix, whose strain energy density function is given by

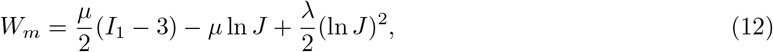

where *µ* is the shear modulus, *λ* is the first Lamé parameter, *I*_1_ is the first invariant of the right Cauchy-Green deformation tensor and *J* is the determinant of the deformation gradient tensor. Fibres are assumed to be incompressible and are modelled using a Neo-Hookean model with a strain energy density function given by

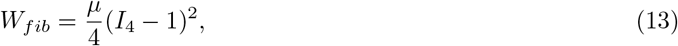

where *I*_4_ is the fourth invariant of the right Cauchy-Green deformation tensor.

#### 2.1.2 Geometry, Initial and Boundary conditions

The geometry used to model the retina is obtained by segmenting the outline of a retinal OCT-B scan (Figure 1a), representing a cross-sectional slice through the macula. We consider one half of this slice, split symmetrically along the fovea, as shown in Figure 1c. The geometry obtained by drawing the outline of the retinal OCT-B scan is 70 *µ*m thick (along Y) at the fovea and 370 *µ*m wide (along X). This 2D profile is extruded in the out-of-plane direction to create a 3D geometry compatible with the FEBio solver, effectively modelling a thin strip of retinal tissue. The aim of our model is to understand the biomechanical effects resulting in oedema as observed in Figure 1b. Hence, the thickness of this strip is set to 1 *µ*m (along Z), which is smaller than the other dimensions of the retina. This choice is based on the assumption that for a thin slice, out-of-plane flow gradients and stresses are negligible, and the in-plane behaviour, which is of primary interest, dominates. This approach is a reasonable simplification for analysing the mechanics in the cross-sectional plane captured by the OCT image. The initial state of the retina is considered to be stress-free, with zero fluid pressure and a uniform baseline solute concentration. The boundary conditions are summarised below and illustrated in Figure 1c.

**Figure 1:**
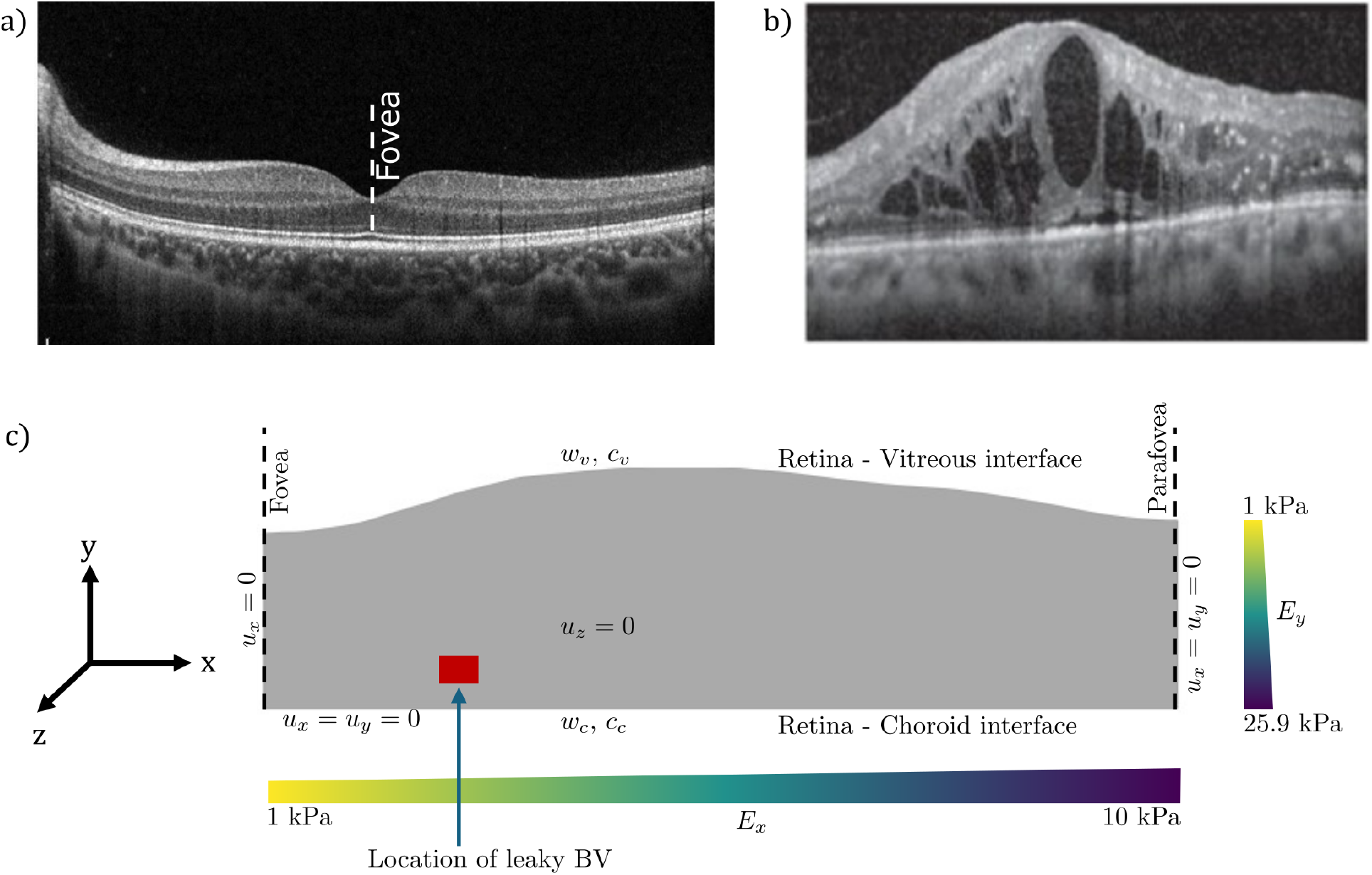
Geometry and Boundary conditions. The OCT-B scan of a healthy retina. Image adapted from [20]. (b) OCT-B scan showing retinal oedema development due to fluid accumulation. Image adapted from [21] (c) Initial geometry of the retina showing the boundary conditions and stiffness map. Stiffness increases from fovea to parafovea, and from vitreous to choroid. The red square indicates the location of leaky blood vessel (BV). Fluid flux boundary conditions (*w*_*v*_, *w*_*c*_). Displacement boundary conditions (*u*_*x*_, *u*_*y*_, *u*_*z*_). Solute concentration boundary conditions (*c*_*v*_, *c*_*c*_).

##### Displacement Boundary Conditions

To simulate the mechanical constraints on the tissue, the retina-choroid interface at the bottom of the model is fully fixed, preventing any movement (*u*_*x*_ = *u*_*y*_ = 0). The right boundary representing the side that is away from the fovea is also fully fixed. The left boundary, representing the foveal axis of symmetry, is constrained against horizontal movement (*u*_*x*_ = 0). This boundary condition simulates the mechanical constraint imposed by the surrounding retinal tissue, which restricts lateral expansion while allowing vertical deformation due to fluid accumulation. Further, the retina is constrained against out-of-plane movement (*u*_*z*_ = 0).

##### Fluid and Solute Boundary Conditions

Fluid and solute exchange occurs at the top (vitreous-retinal) and bottom (retina-choroid) interfaces. Fluid flux (*w*) across these semi-permeable membranes is governed by Starling’s law:

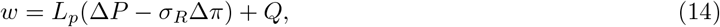

where *L*_*p*_ is the hydraulic conductivity of the interface, Δ*P* is the hydrostatic pressure difference across the interface, *σ*_*R*_ is the reflection coefficient of the interface, Δ*π* is the osmotic pressure difference across the interface and *Q* is the fluid source term. Hydraulic conductivity (*L*_*p*_) signifies how permeable the interface is. Although asymmetric permeability has been observed at the vitreous-retinal interface [19], in this article, we assume equal permeability for flow in both directions. Further, we prescribe the solute concentration at the retina-vitreous interface (*c*_*v*_) and retina-choroid interface (*c*_*c*_) as 0.011 mM and 0.053 mM, respectively. These values represent the typical concentrations of albumin in the vitreous and choroid [6]. Following the FEBio convention, if fluid and solute boundary conditions are not specified, respective fluxes are prescribed as zero.

##### Pathological vascular leakage

Pathological vascular leakage is modelled by introducing a fluid source at a single location within the retinal domain, as shown in Figure 1c. This localised source is intended to represent the aggregate effect of microvascular leakage from the deeper capillary plexus, projected onto the 2D cross-sectional plane of the OCT image, rather than a single ruptured vessel. While this is a simplification, it captures the essential effect of increased fluid influx into the retinal tissue due to vascular leakage. The magnitude of this source term is varied systematically to study its effect on oedema development.

#### 2.1.3 Model parameters and numerical implementation

The retina is a heterogeneous material with stiffness varying along the length and width of the retina [5, 6]. This is mainly due to the varying density and orientation of the fibres in the retinal ECM. Permeability and diffusivity also vary spatially due to the heterogeneity in cell types and ECM composition [6]. We implement heterogeneity in the stiffness as shown in Figure 1c. Similarly, diffusion and permeability are assumed to vary linearly and inversely with stiffness. This is based on the physical principle that a denser ECM (which corresponds to higher stiffness) presents greater resistance to fluid and solute movement, thereby reducing permeability and diffusivity. Additionally, it is also seen that stiffness is higher near arteries and veins [22], which we neglect in this study. The parameters used in the model are summarised in Table 1. Any variations to the baseline parameters are mentioned in the respective sections.

**Table 1:** Model parameters used in this article.

| Parameter | Symbol | Value | Reference |
| --- | --- | --- | --- |
| Young's modulus of retinal ECM | $E_x$ | 1-10 kPa | [23] |
| Young's modulus of retinal ECM | $E_y$ | 1-25.9 kPa | [24] |
| Permeability | k | 1e2 - 1e3 $\mu\text{m}^4/(\text{nNs})$ | Estimated, [6] |
| Diffusivity of albumin | D | 1-10 $\mu\text{m}^2/\text{s}$ | [6] |
| Osmotic coefficient | $\Phi$ | 1 | Assumed |
| Free diffusivity of albumin | $D_0$ | 100 $\mu\text{m}^2/\text{s}$ | [6] |
| Solubility of albumin | $\tilde{k}$ | 1 | [6] |
| Hydraulic conductivity of ILM | $L_p^I$ | 0.01 $\mu\text{m}/(\text{s kPa})$ | [6] |
| Reflection coefficient of ILM | $\sigma$ | 1 | [6] |
| Hydraulic conductivity of RPE | $L_p^R$ | 0 $\mu\text{m}/(\text{s kPa})$ | [6] |
| Reflection coefficient of RPE | $\sigma_R$ | 1 | [6] |
| Fluid pressure in vitreous | $P_{\text{vitreous}}$ | 1 kPa | [25] |
| Fluid pressure in choroid | $P_{\text{choroid}}$ | 2 kPa | Estimated, [26] |
| Universal gas constant | R | 8.314 J/(mol K) | - |
| Temperature | $\theta$ | 310 K | - |
| Vitreous solute concentration | $c_v$ | 0.011 mM | [6] |
| Choroid solute concentration | $c_c$ | 0.053 mM | [6] |
| Diffusivity of anti-VEGF | $D_{\text{anti-VEGF}}$ | 2 $\mu\text{m}^2/\text{s}$ | Estimated, [27] |
| Free Diffusivity of anti-VEGF | $D_{0,\text{anti-VEGF}}$ | 10 $\mu\text{m}^2/\text{s}$ | Assumed |
| Solubility of anti-VEGF | $\tilde{k}_{\text{anti-VEGF}}$ | 0.001 | Assumed |
| Diffusivity of VEGF | $D_{\text{VEGF}}$ | 0.001 $\mu\text{m}^2/\text{s}$ | Assumed |
| Free Diffusivity of VEGF | $D_{0,\text{VEGF}}$ | 1 $\mu\text{m}^2/\text{s}$ | Assumed |
| Solubility of VEGF | $\tilde{k}_{\text{VEGF}}$ | 0.001 | Assumed |
| Diffusivity of anti-VEGF-VEGF complex | $D_{\text{anti-VEGF-VEGFcomplex}}$ | 0.001 $\mu\text{m}^2/\text{s}$ | Assumed |
| Free Diffusivity of anti-VEGF-VEGF complex | $D_{0,\text{anti-VEGF-VEGFcomplex}}$ | 1 $\mu\text{m}^2/\text{s}$ | Assumed |
| Solubility of anti-VEGF-VEGF complex | $\tilde{k}_{\text{anti-VEGF-VEGFcomplex}}$ | 0.001 | Assumed |
| Solid volume fraction | $\phi_s$ | 0.2-0.4 | [6] |
| Young's modulus of blood vessel | $E_{bv}$ | 10 kPa | [23] |
| Permeability of blood vessel to water | $k_{bv}$ | 100 $\mu\text{m}^4/(\text{nNs})$ | Estimated, [6] |
| Osmotic coefficient of blood vessel | $\Phi_{bv}$ | 1 | Assumed |
| Rate constant of anti-VEGF-VEGF binding | $k_{on}$ | 150 $\mu\text{m}^3/(\text{mol s})$ | [15] |
| Reference solute concentration | $c_{ref}$ | 0.0026 mol/$\mu\text{m}^3$ | Assumed |

The model parameters used in this study were sourced from a combination of existing literature and phenomenological estimation. Where available, parameter values were adopted directly from experimental or previous modelling studies as cited in Table 1. For parameters not readily available in the literature, values were estimated to ensure that the model’s output qualitatively matched established physiological and pathological observations. Given the qualitative nature of the validation in this initial study, a systematic sensitivity analysis to fully explore the parameter space was considered beyond the current scope. Such an analysis would, however, be a valuable next step to quantify parameter uncertainty and identifiability. In the cases when we consider fibres in the retina, we assume that fibre orientation is either z-shaped or vertical. Accordingly, the fibre orientation is assumed to affect permeability and diffusion tensors.

The governing equations with the chosen material parameters are implemented in FEBio [18] using the multiphasic material model. The geometry is meshed using quad elements with a non-uniform mesh size, with a finer mesh near the macula. To reduce computational time during parameter sweeps, the restart option in FEBio is used to continue simulations from a previously saved state. Finally, we define macular displacement as the vertical displacement (along Y) of the node located at the centre of the fovea, which corresponds to the bottom of the foveal pit. A positive displacement indicates an increase in retinal thickness (oedema), while a negative displacement signifies retinal thinning.

A note on the interpretation of time and the quasi-static approach is warranted to clarify the source of time-dependence in our model. At each step of the simulation, we solve for the mechanical equilibrium of the retinal tissue under a given set of loads and boundary conditions, as shown in the momentum balance equation (Equation 6). Further, the time-dependence arises from the transient nature of solute transport and fluid flow. The simulation progresses through a sequence of these mechanical equilibrium states. The ‘time’ in our simulation is the parameter that tracks this progression. For example, in our anti-VEGF simulations, the concentration of the drug changes over time due to diffusion. This change in concentration then modifies the vascular leakage rate, which acts as a time-varying load on the tissue. The tissue then deforms to find a new mechanical equilibrium in response to this new load. Therefore, the overall process is dynamic, but solved using a quasi-static approach where the mechanics are assumed to equilibrate instantly to changes in the fluid and solute environment. While our simulations reach equilibrium over computational timescales of minutes, the corresponding physiological processes of oedema development or resolution occur over months to years. This approach is a deliberate simplification, enabling a computationally tractable investigation into the fundamental mechanistic relationships between pressure, flow, and deformation, rather than a precise chronological prediction of the disease and treatment course.

## 3 Results

### 3.1 RPE pumping keeps the retina dry

It is well established that in addition to acting as a barrier at the retina-choroid interface, the RPE pumps fluid out of the retina and into the choroid [28]. Since the hydrostatic pressure of the choroid is higher than that of the retina, an active pumping mechanism is essential for maintaining physiological retinal thickness [25]. Accordingly, to establish a physiological baseline for our model, we first simulated the role of active pumping by the RPE in maintaining retinal homeostasis. This baseline case serves as a reference against which we compare the pathological conditions simulated in subsequent sections. We modelled the physiological RPE as a boundary with zero hydraulic conductivity, representing an impermeable retina-choroid barrier, and included a non-zero fluid source term (Q) to signify active pumping by the RPE (Equation 14). This approach models fluid transport across the RPE-choroid interface as a purely transcellular process, where fluid is actively pumped through the RPE cells, assuming the paracellular pathway via tight junctions is closed. To explore the balance between active pumping and passive fluid intake due to permeable RPE, we systematically varied both the RPE pumping rate and the hydraulic conductivity of the RPE barrier (Figure 2a).

**Figure 2:**
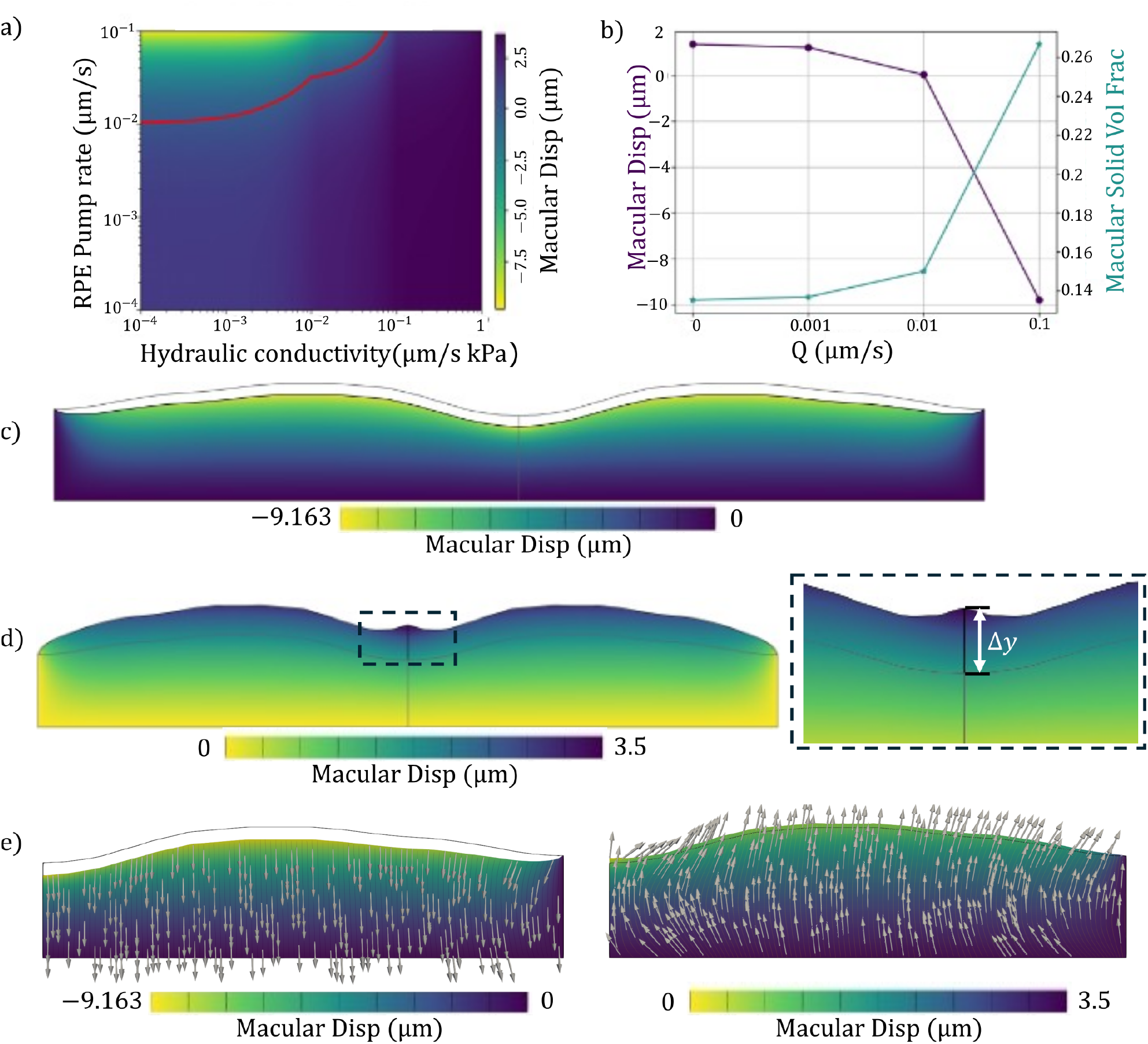
(a) Retinal displacement as a function of RPE pumping rate and hydraulic conductivity, with a contour line marking zero displacement. (b) Macular thickness reduces with increasing RPE pumping rate (left y-axis) while solid volume fraction increases (right y-axis). Hydraulic conductivity is set to 0. (c) Retinal thickness decreases due to active pumping by RPE (grey line indicates the outline of undeformed retina). (d) Retinal thickness increases due to loss of active pumping by RPE. The macular thickness is enlarged in the box. (e)Fluid flux with active pumping by RPE (left) and without active pumping by RPE (right). Arrows indicate fluid flux direction.

Our simulations show that the macular displacement is governed by the interplay between active fluid removal by the RPE pump and passive fluid influx due to the barrier’s leakiness (hydraulic conductivity). For any given pump rate, an increase in hydraulic conductivity leads to greater fluid accumulation and retinal swelling. Conversely, for a given hydraulic conductivity (level of leakiness of the retina-choroid barrier), a higher pumping rate results in a thinner retina (Figure 2a). The red contour line in (Figure 2a) delineates the critical threshold where these two processes are in balance, resulting in zero net displacement. This contour separates the physiological state (net fluid removal) from the pathological state of oedema (net fluid accumulation), highlighting the non-linear coupling between RPE pumping and permeability in maintaining retinal fluid homeostasis.

Further, keeping the hydraulic conductivity constant at zero, we varied the RPE pumping rate (Q) from 0 to 0.1 *µ*m/s to explore its effect on retinal thickness and solid volume fraction (Figure 2b). Earlier studies report RPE fluid transport rates in the range of 0.011-0.028 *µ*m/s [29, 14]. Our model showed that retinal thickness decreases for Q ≥ 0.01 *µ*m/s, which falls within the reported physiological range. Furthermore, we measured the solid volume fraction in the macular region of the retina for each case and found that it increases with the RPE pumping rate, indicating a drier retina (Figure 2b). We observed that this increase is most prominent in the macular region, highlighting the critical role of RPE pumping in keeping this area dry (Figure S1). Conversely, when the RPE fails to pump fluid (Q = 0), fluid accumulates, leading to a decreased solid volume fraction, which signifies the development of oedema (Figure S1).

Accordingly our simulations show that the retinal thickness (measured as the displacement at the tip of the macula) reduces in the physiological case (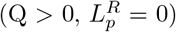) (Figure 2c) and increases in the pathological case (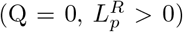) (Figure 2d). We further observe that in the physiological case, fluid flows from the vitreous to the choroid (Figure 2e, left), while in the pathological case, fluid flux reverses direction and flows from choroid towards vitreous (Figure 2e, right). These findings are consistent with experimental observations that carbonic anhydrase inhibitors, such as acetazolamide, can reduce oedema by stimulating active RPE pumping [30].

### 3.2 BV leakage along with dysfunctional RPE aggravates oedema development

Next, we investigated the combined effect of blood vessel (BV) leakage and RPE dysfunction, which often coexist in retinal pathologies such as diabetic macular oedema [3]. To simulate this, we introduced a fluid source term that represents a leaky blood vessel, allowing fluid to enter retinal tissue. We then simulated different conditions by varying both the prescribed rate of BV leakage and RPE active pumping. Keeping the RPE pumping rate constant at 0.1 *µ*m/s, we observed that increasing BV leakage leads to a significant increase in retinal thickness and a corresponding decrease in solid volume fraction, indicating oedema development (Figure 3c). We also observe that the fluid accumulates and retinal thickness increases most prominently in the macular region.

**Figure 3:**
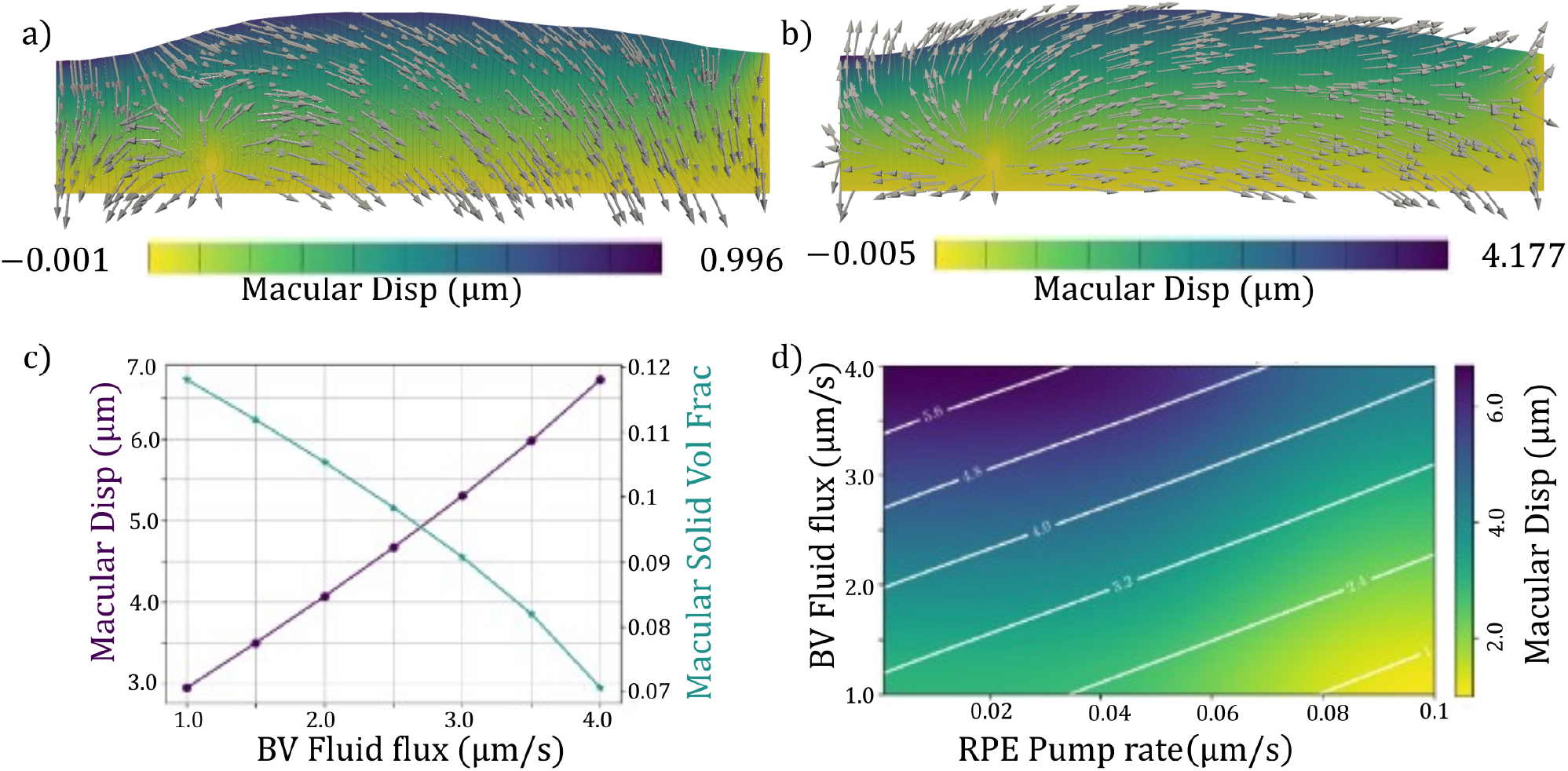
(a) Retinal thickness with a BV leakage of 1 *µ*m/s. Arrows indicate the direction of fluid flux. (b) Retinal thickness with a BV leakage of 4 *µ*m/s. Arrows indicate the direction of fluid flux.(c) Macular thickness increases with increasing BV leakage (left y-axis) while macular solid volume fraction decreases (right y-axis). (d) Retinal displacement as a function of RPE pumping rate and BV leakage, with contour lines drawn at different thicknesses.

Further, our simulations showed that the direction of fluid flux depends on the magnitude of BV leakage. When the BV leakage was set to 1 *µ*m/s, the fluid flows from vitreous towards choroid, as in the physiological case (Figure 3a). However, when the BV leakage is increased to 4 *µ*m/s, the fluid flow reverses direction, moving from the leaky blood vessel towards both the vitreous and choroid (Figure 3b). This reversal in fluid flow is indicative of the pathological state of oedema, which is a combination of the magnitude of BV fluid flux and RPE pump rate.

Further, our model predicts that BV leakage and RPE pump rate synergistically worsen oedema development. As BV leakage increases and RPE pumping decreases, the retina accumulates more fluid, leading to a significant increase in thickness at the macula and a corresponding decrease in the solid volume fraction, indicating a wetter tissue state. Plotting the heatmap of the macular displacement as a function of RPE pump rate and BV leakage, with hydraulic conductivity set at 0.01 *µ*m/s/kPa we observe that the contours are linear (Figure 3d). This indicates that the macular displacement is a function of both RPE pump rate and BV leakage. These findings underscore the complex interplay between vascular leakage, RPE function, and hydraulic permeability in the pathogenesis of retinal oedema. Further, we estimate that including active cellular responses affecting the RPE pump rate and BV leakage, the interaction would become non-linear.

### 3.3 Anti-VEGF treatment restores retinal thickness in a concentration-dependent manner

Next, we modelled the effect of anti-VEGF therapy, the first-line treatment in many retinal diseases [3]. Pathological fluid accumulation is often driven by VEGF, which disrupts the blood-retinal barrier and increases vascular permeability [31]. Patients experiencing macular oedema are commonly treated with intravitreal injections of anti-VEGF agents, which bind to VEGF and inhibit its activity, thereby reducing vascular leakage and promoting fluid resorption [3]. Decades of clinical studies have shown that patients’ response to anti-VEGF treatment varies with time and the drug used [32]. However, the reason behind this variation is not yet understood. To simulate this treatment, we introduced an anti-VEGF solute that diffuses within the retina. We initially allow the thickness of the retina to increase upto 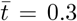 (1/3 of the total simulation time). Then we introduce an anti-VEGF solute at the vitreous-retina interface which is expected to diffuse towards the location of the leaky blood vessel. For these simulations, all other material and transport parameters, such as tissue stiffness, permeability, and diffusion coefficients for albumin, were maintained at their baseline values as detailed in Table 1. Following [15], we further model the binding between anti-VEGF solutes and free VEGF molecules to form the anti-VEGF-VEGF complex following first order chemical kinetics (Equation 15). We assumed that the prescribed fluid leakage from blood vessels (BV fluid flux) decays exponentially with the concentration of anti-VEGF-VEGF complex (Equation 16). (BV fluid flux)_*t*_ is the prescribed fluid flux at the location of leaky blood vessel at a time *t*, (BV fluid flux)_0_ is the initial prescribed fluid flux, and *C*_*ref*_ is a constant reference solute concentration. The choice of an exponential decay function is qualitatively inspired by clinical data, such as the DRCR.net Protocol T study [32], and has been discussed in the Discussion section. While in this study, we set *C*_*ref*_ based on the phenomenological observation, in future work, this can be modified to match the clinical studies. Accordingly, our simulations show that anti-VEGF solutes reduce fluid leakage, leading to a decrease in retinal thickness. We observed that higher anti-VEGF concentrations lead to a more rapid restoration of retinal thickness to pre-oedema levels after its injection at 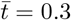 (Figure 4c). While our model shows that a single injection is sufficient to restore retinal thickness, in real clinical scenarios, the patient is subjected to repeated anti-VEGF injections at regular intervals to maintain desired retinal thickness and visual acuity [32]. The discrepancy between our model’s prediction of a single-dose resolution and the clinical need for repeated injections stems from key simplifications. Primarily, we model vascular leakage as a single point source, thereby neglecting the complex, distributed network of leaky vessels often seen in DMO. Furthermore, the model omits the physiological mechanisms of anti-VEGF drug clearance and breakdown, which are critical for determining long-term efficacy. These simplifications were intentionally made to isolate the core mechanical and transport interactions. The underlying porous medium framework is, however, capable of incorporating these more realistic features in future work, where we will incorporate such complexity and study the need for repeated dosing to better mimic clinical treatment regimens.

**Figure 4:**
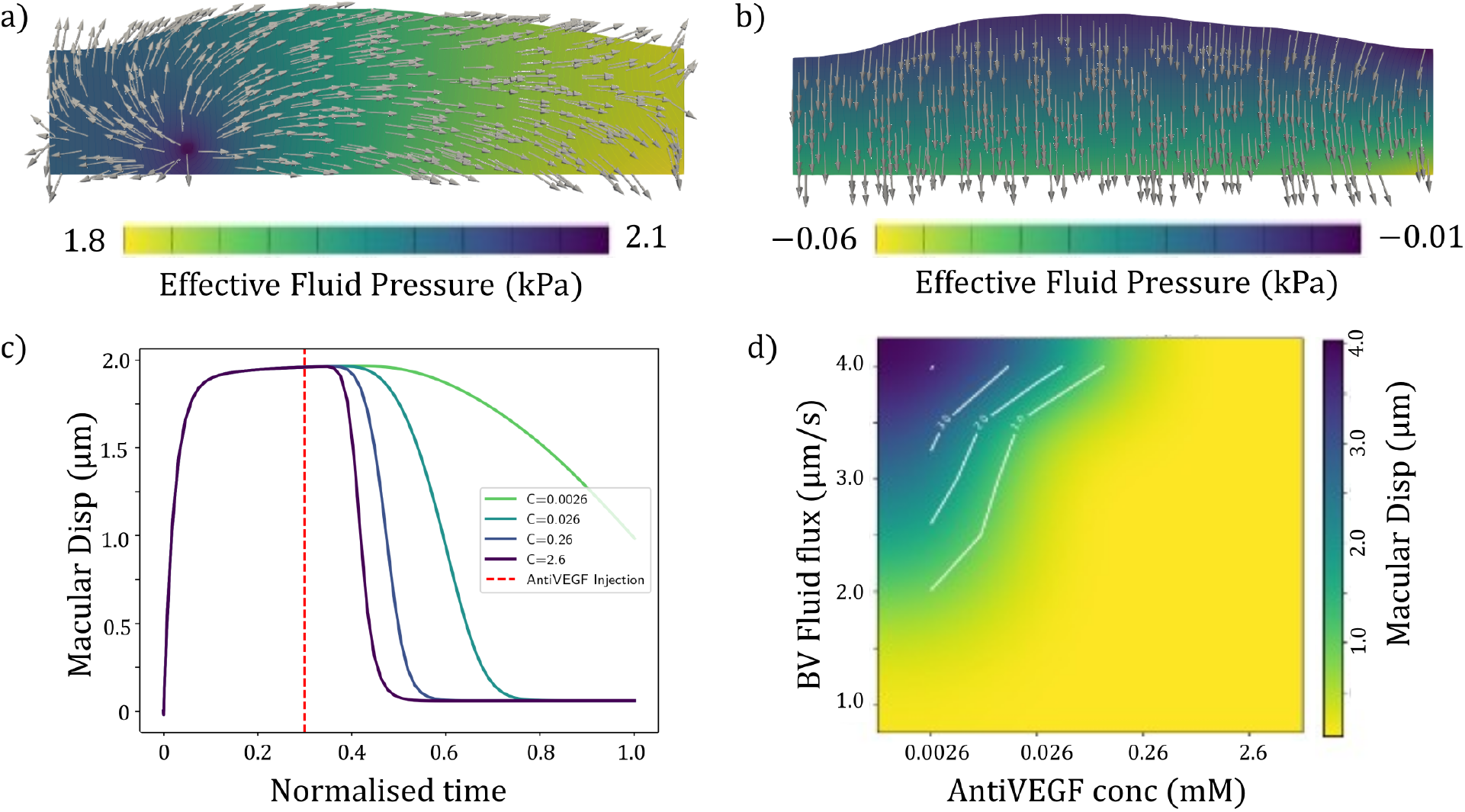
(a) Fluid pressure in the retina increases with BV leakage. The arrows indicate the direction of fluid flux at the steady state. Retinal thickness also increases. (b) Fluid pressure reduces due to anti-VEGF treatment. Arrows indicate the direction of fluid flux at the steady state. Retinal thickness also reaches the pre-oedema levels. (c) Retinal thickness over time after anti-VEGF treatment at different concentrations (d) Retinal displacement as a function of anti-VEGF concentration and BV leakage, with contour lines marking different thicknesses.

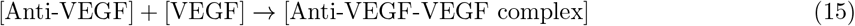

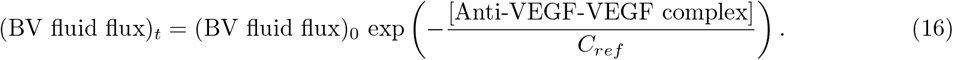

Further, we studied the coupling between BV leakage and anti-VEGF concentration on retinal thickness. We observed that as the BV leakage increases, a higher anti-VEGF concentration is required to restore retinal thickness to pre-oedema levels (Figure 4d). We also observed that the contour lines are non-linear due to the second-order diffusion equation governing anti-VEGF transport. This implies that the required drug dosage does not scale linearly with the severity of vascular leakage, potentially explaining why standard dosing regimens may fail in severe cases. While these results align with clinical observations that anti-VEGF treatments significantly reduce macular oedema by stabilising the blood-retinal barrier and promoting fluid resorption, further work is needed to match the timescales of treatment and response. Additionally, we observed that the fluid flux patterns within the retina change with anti-VEGF concentration. At low anti-VEGF concentrations, fluid flux is directed from the leaky blood vessel towards both the vitreous and choroid, along with positive fluid pressure indicative of oedema (Figure 4a). However, at high anti-VEGF concentrations, the blood vessel leakiness is stopped, and the fluid flux direction is from the vitreous towards the choroid, along with negative fluid pressure, indicating a resolved physiological state (Figure 4b). These findings suggest that anti-VEGF therapy not only reduces retinal thickness but also restores normal fluid dynamics within the retina.

### 3.4 Z-shaped Müller cell orientation protects against oedema development but hinders anti-VEGF delivery

Finally, we explored the influence of the orientation of Müller cells on retinal mechanics and oedema development. The Müller cells regulate the mechanical environment of the retina, including providing structural support [6] and regulating fluid and solute transport [3]. In a healthy retina, Müller cells form a Z-shaped distribution and in pathological conditions such as retinal detachment and macular oedema, the Müller cells progressively straighten [7], as shown in the schematic (Figure 5c). Accordingly, we modelled the retina with fibres and varied their orientation to assess their impact on retinal thickness and fluid dynamics. We consider two cases: (1) fibres aligned in the physiological Z-shaped configuration, and (2) fibres oriented vertically, representing the pathological state (Figure 5a,b). Although perfect vertical alignment is an idealisation not strictly observed *in vivo*, we analyse this limiting case to elucidate the impact of fibre orientation on retinal biomechanics. Green lines indicate vertical fibres (perpendicular to RPE surface) while red lines indicate non-vertical fibres (Z-shaped). Further, the diffusivity and permeability are assumed to be dependent on the orientation of the fibre as shown in equations

**Figure 5:**
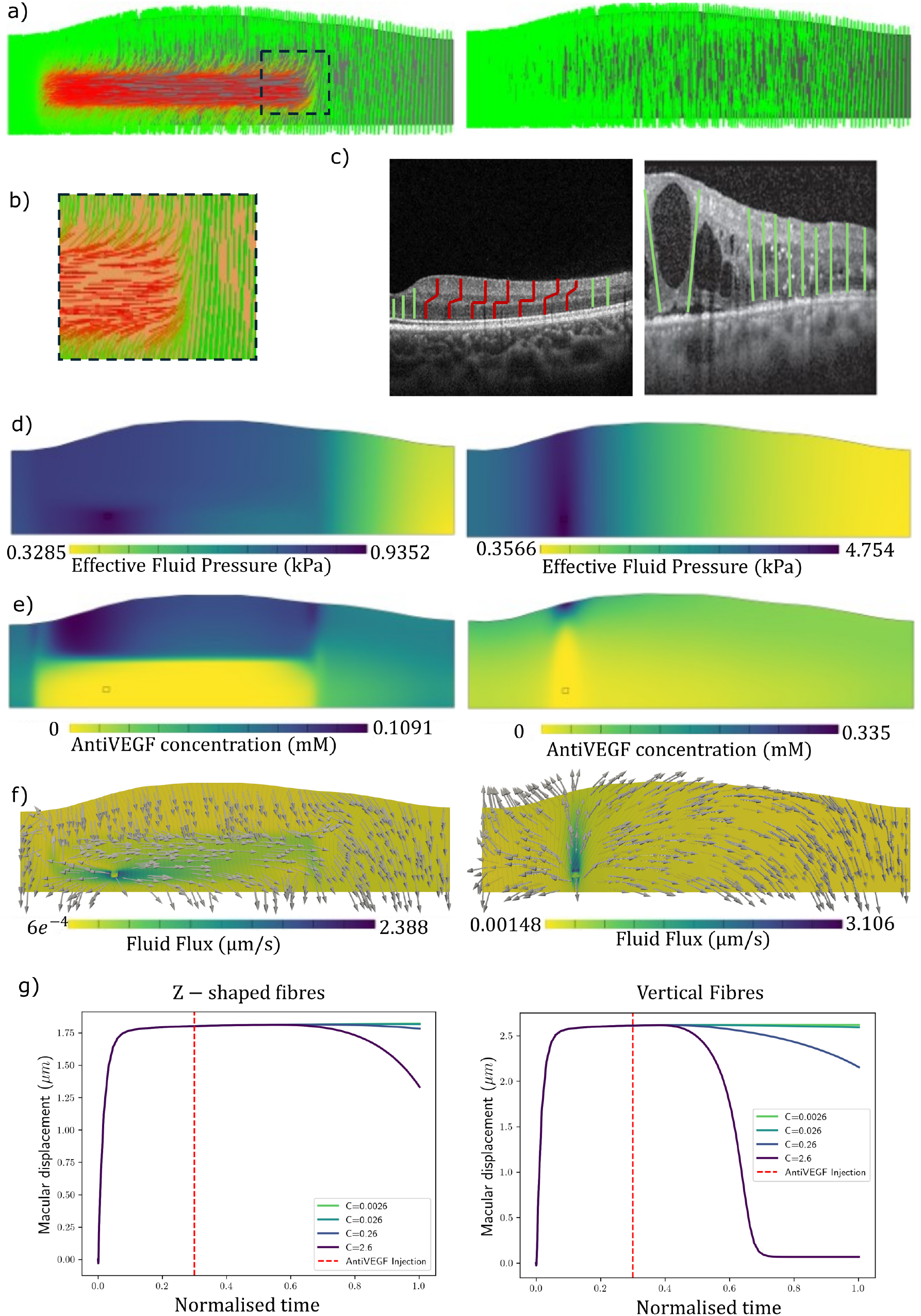
(a) Fibre orientation in z-shaped (left) and vertical (right) configurations. Green indicates vertical fibre and red indicates non-vertical fibres.(b) zoomed-in view of fibre orientation in z-shaped (left) configuration. (c) Schematic of fibre orientation in z-shaped (left) and vertical (right) configurations overlayed on physiological [20] and pathological [21] OCT images, respectively. (d) Effective fluid pressure distribution in z-shaped (left) and vertical (right) fibre configurations. Square drawn within the retina indicates the region of leaky BV. (e) Anti-VEGF concentration distribution in z-shaped (left) and vertical (right) fibre configurations. Square drawn within the retina indicates the region of leaky BV. (f) Fluid flux distribution in z-shaped (left) and vertical (right) fibre configurations. Square drawn within the retina indicates the region of leaky BV. (g) Retinal thickness over time after anti-VEGF treatment in z-shaped (left) and vertical (right) fibre configurations.

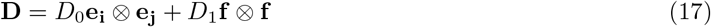

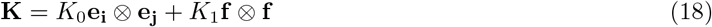

where **D** is the diffusion tensor, **K** is the permeability tensor, *D*_0_ and *K*_0_ are the isotropic diffusivity and permeability, *D*_1_ and *K*_1_ are the anisotropic diffusivity and permeability along the fibre direction, **f** is the unit vector along the fibre direction, and **e**_**i**_ and **e**_**j**_ are the unit vectors along the coordinate axes.

Our simulations revealed that fibre orientation significantly modulates the mechanical response of the retina to BV leakage. For a given level of vascular leakage, models with vertically oriented fibres exhibited higher maximum effective fluid pressure (Figure 5d, right) compared to those with the physiological Z-shaped configuration (Figure 5d, left). This resulted in greater macular thickness in the vertical fibre case (Figure 5g) and a lower solid volume fraction, indicating a wetter retina. The Z-shaped fibre configuration was observed to impede fluid and solute flow towards the macular region when compared to the vertical configuration (Figure 5e,f). Similar to the earlier problem, we introduced the anti-VEGF drug at 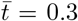 and found that the anti-VEGF concentration was low near the leaky blood vessel in the case of Z-shaped fibres compared to vertical fibres (Figure 5e). Similarly, the fluid flux was also lower in the case of Z-shaped fibres compared to vertical fibres (Figure 5f). Further, we observed that the time taken for the retinal thickness to reduce to pre-oedema levels after anti-VEGF treatment was lower in the case of vertical fibres (Figure 5g, right) compared to Z-shaped fibres (Figure 5g, left), even though thickness was higher. These findings suggest that the orientation of Müller cells plays a crucial role not only in modulating the retina’s biomechanical response to oedema, but also its recovery following treatment. We further studied the impact of diffusivity and permeability anisotropy on retinal thickness and anti-VEGF delivery (Figure S2). We observed that anisotropy in permeability has a higher influence on anti-VEGF efficacy than anisotropy in diffusivity. However, its clinical relevance needs to be further explored.

## 4 Discussion and Conclusion

In this article, we developed a multiphysics computational model of the retina that integrates anatomically inspired geometry, vascular leakage, retinal biomechanics, fibre distribution morphology, fluid dynamics, and solute transport to study oedema development and anti-VEGF therapy response. Our model successfully captures key physiological and pathological phenomena observed in retinal diseases such as diabetic macular oedema (DMO). We demonstrated that the mechanism of active pumping by the retinal pigment epithelium (RPE) was essential for recapitulating the physiological retinal thickness. The model showed that the impaired pumping led to fluid accumulation and oedema. This finding aligns with experimental observations that enhancing RPE function can mitigate oedema [30]. Further, we showed that blood vessel leakage and RPE dysfunction synergistically exacerbate oedema development, highlighting the complex interplay between vascular and epithelial factors in retinal fluid homeostasis. Our simulations also revealed that anti-VEGF therapy effectively reduces retinal thickness in a concentration-dependent manner, consistent with clinical outcomes. A key insight from our model is the dual role of Müller cell architecture in both protecting against and complicating the treatment of macular oedema. Our simulations demonstrate that the physiological Z-shaped orientation of these cells provides a biomechanical advantage, stiffening the retina and resisting fluid accumulation. However, this protective structure simultaneously acts as a barrier, impeding the diffusion of anti-VEGF drugs to leaky vessels. This finding reveals a critical trade-off between the retina’s structural integrity and its response to therapy, offering a potential explanation for the variable efficacy of anti-VEGF treatments observed clinically. It highlights that cellular morphology is a critical determinant of both disease progression and therapeutic outcome.

While multiphysics modelling is a vibrant and rapidly advancing field, its translation into routine clinical practice remains a significant hurdle across many areas of medicine, not just ophthalmology [13, 5]. This ‘bench-to-bedside’ gap highlights a broad, well-recognised challenge. A key bottleneck in this process, and a significant challenge for the model presented here, is the robust parameterisation of the underlying biological processes. Our model incorporates numerous parameters (Table 1), many of which are difficult to measure directly *in vivo* and exhibit considerable variability in the literature. While we obtained many parameters from established experimental studies, others were estimated or assumed based on prior modelling work or qualitative observations. This introduces uncertainty into the model predictions, which must be carefully considered when interpreting the results. To rigorously assess the impact of this uncertainty, a systematic sensitivity analysis would be required, which is outside the scope of this article. Such an analysis would identify the most influential parameters, thereby guiding future experimental research to focus on obtaining more precise measurements. Furthermore, this would be a critical step towards clinical translation, where techniques like Bayesian inference could be used to calibrate the model against patient-specific clinical data (e.g., longitudinal OCT measurements), allowing for the inference of patient-specific parameters. These inferred parameters could themselves serve as novel biomarkers, offering deeper insights into an individual’s disease state.

### Phenomenological model of anti-VEGF efficacy

Equation 16 describes a phenomenological relationship intended to capture the dose-dependent reduction in vascular leakage following anti-VEGF administration. The choice of an exponential decay function is qualitatively inspired by clinical data, such as the DRCR.net Protocol T study, which demonstrated that anti-VEGF agents lead to a rapid initial reduction in macular thickness (a surrogate for leakage), with the effect tapering over time (Figure S4) [32]. This functional form models the saturating effect of the drug, where initial increases in concentration have a substantial impact on reducing leakage, while at higher concentrations, the effect plateaus. Hence, we use an exponential relation noting that the current study is not designed for quantitative dosing predictions but rather to investigate mechanistic trends. Further, the concentration scaling parameter *C*_*ref*_ is a phenomenological constant that modulates drug efficacy, and its calibration would require extensive clinical data, which is beyond the scope of the current study.

Finally, our model has several limitations that warrant discussion. First, our model is based on a 2D crosssection of the retina, extruded to a small thickness. This simplification assumes that out-of-plane stresses and flow gradients are negligible. While this is a reasonable starting point for understanding the primary mechanics in the OCT plane, a full 3D model of the curved retinal geometry would be necessary to capture more complex, three-dimensional fluid flows and mechanical interactions. Future work could include a sensitivity analysis to slice thickness and, ultimately, extension to a full 3D representation. Second, while we incorporated key biomechanical and transport phenomena, we did not account for active cellular responses, including cell degeneration, that are commonly observed in retinal pathologies. As a result, our model predicted that with an increase in IOP, the retinal thickness increases (Figure S3), which is in contrast to clinical observations where increased IOP often leads to retinal thinning due to ischaemia-induced cell loss [33]. Future work should integrate cellular dynamics and degeneration processes to more accurately capture disease progression. Third, we simplified the vascular network to a single location indicating a leaky blood vessel, whereas in reality, the retina contains a complex vasculature with multiple leakage sites. Expanding the model to include a more detailed vascular architecture would provide a more comprehensive understanding of oedema development. Finally, our simulations focused on short-term responses to anti-VEGF therapy following a single injection and qualitative comparison with clinical observations. In clinical practice, patients often receive repeated treatments over extended periods. Future studies should incorporate longitudinal treatment regimens to better mimic clinical scenarios and assess long-term therapeutic outcomes. Furthermore, the current continuum approach could be refined to capture structural biomarkers such as the disorganisation of retinal inner layers (DRIL) [34], which correlates with visual outcomes in DMO. Additionally, the model’s capacity to predict fluid pressure accumulation offers a pathway to simulate the mechanical onset of retinal detachment, where hydrostatic forces overcome the adhesion at the sub-retinal space.

Thus, our porous media-based mathematical model provides a comprehensive framework for understanding the multifaceted mechanisms driving retinal oedema and evaluating therapeutic interventions. The insights obtained from this model enhance our understanding of retinal pathophysiology and may inform the development of personalised treatment strategies for retinal diseases such as diabetic macular oedema.

## 5 Code availability

FEBio input files and scripts used for simulations are available as open source at: https://github.com/bkprdp/Diabetic-Macular-Oedema-development-and-therapy.git

## 6 Acknowledgements

RS and SWS would like to acknowledge funding from Cancer Research UK ((28276 and C44767/A29458)) and EPSRC ((EP/W007096/1)).

The authors acknowledge the use of Google’s Gemini (2.5 Pro) for assistance with manuscript preparation. Disclosure: PK: None, EB: None, PL: None, RS: None, SWS: None.

## Notes

### Competing Interest Statement

The authors have declared no competing interest.

## References

[1] Ning Cheung, Chiu Ming Gemmy Cheung, Steven James Talks, and Tien Yin Wong. Management of diabetic macular oedema: new insights and global implications of DRCR protocol V. Eye, 34(6):999–1002, June 2020.

[2] Zhen Ling Teo, Yih-Chung Tham, Marco Yu, Miao Li Chee, Tyler Hyungtaek Rim, Ning Cheung, Mukharram M. Bikbov, Ya Xing Wang, Yating Tang, Yi Lu, Ian Y. Wong, Daniel Shu Wei Ting, Gavin Siew Wei Tan, Jost B. Jonas, Charumathi Sabanayagam, Tien Yin Wong, and Ching-Yu Cheng. Global Prevalence of Diabetic Retinopathy and Projection of Burden through 2045: Systematic Review and Meta-analysis. Ophthalmology, 128(11):1580–1591, November 2021.

[3] Jingfa Zhang, Jingxiang Zhang, Chaoyang Zhang, Jingting Zhang, Limin Gu, Dawei Luo, and Qinghua Qiu. Diabetic Macular Edema: Current Understanding, Molecular Mechanisms and Therapeutic Implications. Cells, 11(21):3362, January 2022.

[4] Henry A Resnikoff, Charles G Miller, and Jean E Schwarzbauer. Implications of fibrotic extracellular matrix in diabetic retinopathy. Exp Biol Med (Maywood), 247(13):1093–1102, July 2022.

[5] Laura Prieto-López, Xandra Pereiro, and Elena Vecino. The mechanics of the retina: Müller glia role on retinal extracellular matrix and modelling. Front. Med., 11, September 2024.

[6] Alessia Ruffini, Mariia Dvoriashyna, Andrea Govetto, Mario R. Romano, and Rodolfo Repetto. A Mathematical Model of Interstitial Fluid Flow and Retinal Tissue Deformation in Macular Edema. Invest. Ophthalmol. Vis. Sci., 65(11):19, September 2024.

[7] Andrea Govetto, Jean-Pierre Hubschman, David Sarraf, Marta S. Figueroa, Ferdinando Bottoni, Roberto dell’Omo, Christine A. Curcio, Patrizio Seidenari, Giulia Delledonne, Robert Gunzenhauser, Mariantonia Ferrara, Adrian Au, Gianni Virgili, Antonio Scialdone, Rodolfo Repetto, and Mario R. Romano. The role of Müller cells in tractional macular disorders: an optical coherence tomography study and physical model of mechanical force transmission. British Journal of Ophthalmology, 104(4):466–472, April 2020.

[8] Laila Aldokhail, Abdulaziz Alhadlaq, Lujain Alaradi, Lamees Alaradi, and Fatimah AlShaikh. Outcomes of Anti-VEGF Therapy in Eyes with Diabetic Macular Edema, Vein Occlusion-Related Macular Edema, and Neovascular Age-Related Macular Degeneration: A Systematic Review. OPTH, Volume 18:3837–3851, December 2024.

[9] Jeffrey De Fauw, Joseph R. Ledsam, Bernardino Romera-Paredes, Stanislav Nikolov, Nenad Tomasev, Sam Blackwell, Harry Askham, Xavier Glorot, Brendan O’Donoghue, Daniel Visentin, George van den Driessche, Balaji Lakshminarayanan, Clemens Meyer, Faith Mackinder, Simon Bouton, Kareem Ayoub, Reena Chopra, Dominic King, Alan Karthikesalingam, Cían O. Hughes, Rosalind Raine, Julian Hughes, Dawn A. Sim, Catherine Egan, Adnan Tufail, Hugh Montgomery, Demis Hassabis, Geraint Rees, Trevor Back, Peng T. Khaw, Mustafa Suleyman, Julien Cornebise, Pearse A. Keane, and Olaf Ronneberger. Clinically applicable deep learning for diagnosis and referral in retinal disease. Nat Med, 24(9):1342–1350, September 2018.

[10] Yukun Zhou, Mark A. Chia, Siegfried K. Wagner, Murat S. Ayhan, Dominic J. Williamson, Robbert R. Struyven, Timing Liu, Moucheng Xu, Mateo G. Lozano, Peter Woodward-Court, Yuka Kihara, Andre Altmann, Aaron Y. Lee, Eric J. Topol, Alastair K. Denniston, Daniel C. Alexander, and Pearse A. Keane. A foundation model for generalizable disease detection from retinal images. Nature, 622(7981):156–163, October 2023.

[11] Ursula Schmidt-Erfurth, Amir Sadeghipour, Bianca S. Gerendas, Sebastian M. Waldstein, and Hrvoje Bogunović. Artificial intelligence in retina. Progress in Retinal and Eye Research, 67:1–29, November 2018.

[12] Emmeline E. Brown, Andrew A. Guy, Natalie A. Holroyd, Paul W. Sweeney, Lucie Gourmet, Hannah Coleman, Claire Walsh, Athina E. Markaki, Rebecca Shipley, Ranjan Rajendram, and Simon Walker-Samuel. Physics-informed deep generative learning for quantitative assessment of the retina. Nat Commun, 15(1):6859, August 2024.

[13] Paul A. Roberts, Eamonn A. Gaffney, Philip J. Luthert, Alexander J. E. Foss, and Helen M. Byrne. Mathematical and computational models of the retina in health, development and disease. Progress in Retinal and Eye Research, 53:48–69, July 2016.

[14] Mariia Dvoriashyna, Alexander J. E. Foss, Eamonn A. Gaffney, and Rodolfo Repetto. Fluid and solute transport across the retinal pigment epithelium: a theoretical model. Journal of The Royal Society Interface, 17(163):20190735, February 2020.

[15] Jessica R. Crawshaw, Eamonn A. Gaffney, Michael Gertz, Philip K. Maini, and Antonello Caruso. Modeling the Ocular Pharmacokinetics and Pharmacodynamics of Ranibizumab for Improved Understanding and Data Collection Strategies in Ocular Diseases. Invest. Ophthalmol. Vis. Sci., 66(6):20, June 2025.

[16] Laurence A. Hutton-Smith, Eamonn A. Gaffney, Helen M. Byrne, Philip K. Maini, Dietmar Schwab, and Norman A. Mazer. A Mechanistic Model of the Intravitreal Pharmacokinetics of Large Molecules and the Pharmacodynamic Suppression of Ocular Vascular Endothelial Growth Factor Levels by Ranibizumab in Patients with Neovascular Age-Related Macular Degeneration. Mol. Pharmaceutics, 13(9):2941–2950, September 2016.

[17] Gerard A. Ateshian, Steve Maas, and Jeffrey A. Weiss. Multiphasic Finite Element Framework for Modeling Hydrated Mixtures With Multiple Neutral and Charged Solutes. Journal of Biomechanical Engineering, 135(11):111001, November 2013.

[18] Steve A. Maas, Benjamin J. Ellis, Gerard A. Ateshian, and Jeffrey A. Weiss. FEBio: Finite Elements for Biomechanics. Journal of Biomechanical Engineering, 134(1):011005, January 2012.

[19] Eva Ramsay, Marja Hagström, Kati-Sisko Vellonen, Susanna Boman, Elisa Toropainen, Eva M. del Amo, Heidi Kidron, Arto Urtti, and Marika Ruponen. Role of retinal pigment epithelium permeability in drug transfer between posterior eye segment and systemic blood circulation. European Journal of Pharmaceutics and Biopharmaceutics, 143:18–23, October 2019.

[20] Inferrera Leandro, Borsatti Lorenzo, Miladinovic Aleksandar, Marangoni Dario, Giglio Rosa, Accardo Agostino, and Tognetto Daniele. OCT-based deep-learning models for the identification of retinal key signs. Sci Rep, 13(1):14628, September 2023.

[21] Abdelhafid Errabih, Mohyeddine Boussarhane, Benayad Nsiri, Abdelalim Sadiq, My Hachem El Yousfi Alaoui, Rachid Oulad Haj Thami, and Brahim Benaji. Identifying Retinal Diseases on OCT Image Based on Deep Learning. International Journal of Online and Biomedical Engineering (iJOE), 18(15):141–159, December 2022.

[22] Kinon Chen and James D. Weiland. Anisotropic and inhomogeneous mechanical characteristics of the retina. Journal of Biomechanics, 43(7):1417–1421, May 2010.

[23] Kinon Chen, Adrian P. Rowley, James D. Weiland, and Mark S. Humayun. Elastic properties of human posterior eye. Journal of Biomedical Materials Research Part A, 102(6):2001–2007, 2014. eprint: https://onlinelibrary.wiley.com/doi/pdf/10.1002/jbm.a.34858.

[24] Yueqiao Qu, Youmin He, Yi Zhang, Teng Ma, Jiang Zhu, Yusi Miao, Cuixia Dai, Mark Humayun, Qifa Zhou, and Zhongping Chen. Quantified elasticity mapping of retinal layers using synchronized acoustic radiation force optical coherence elastography. Biomed. Opt. Express, 9(9):4054, September 2018.

[25] M. F. Marmor. Control of subretinal fluid: experimental and clinical studies. Eye (Lond), 4 (Pt 2):340–344, 1990.

[26] A. Bill, G. Sperber, and K. Ujiie. Physiology of the choroidal vascular bed. Int Ophthalmol, 6(2):101–107, February 1983.

[27] Alessia Ruffini, Alessia Casalucci, Caterina Cara, C. Ross Ethier, and Rodolfo Repetto. Drug Distribution After Intravitreal Injection: A Mathematical Model. Invest. Ophthalmol. Vis. Sci., 65(4):9, April 2024.

[28] Olaf Strauss. The Retinal Pigment Epithelium in Visual Function. Physiological Reviews, 85(3):845–881, July 2005.

[29] B A Hughes, S S Miller, and T E Machen. Effects of cyclic AMP on fluid absorption and ion transport across frog retinal pigment epithelium. Measurements in the open-circuit state. The Journal of general physiology, 83(6):875–899, June 1984.

[30] B. Moldow, Birgit Sander, Michael Larsen, Claus Engler, B. Li, Thomas Rosenberg, and Henrik Lund-Andersen. The effect of acetazolamide on passive and active transport of fluorescein across the blood-retina barrier in retinitis pigmentosa complicated by macular oedema. Graefe’s Arch Clin Exp Ophthalmol, 236(12):881–889, November 1998.

[31] Alejandra Daruich, Alexandre Matet, Alexandre Moulin, Laura Kowalczuk, Michaël Nicolas, Alexandre Sellam, Pierre-Raphaël Rothschild, Samy Omri, Emmanuelle Gélizé, Laurent Jonet, Kimberley Delaunay, Yvonne De Kozak, Marianne Berdugo, Min Zhao, Patricia Crisanti, and Francine Behar-Cohen. Mechanisms of macular edema: Beyond the surface. Progress in Retinal and Eye Research, 63:20–68, March 2018.

[32] Adam R. Glassman, John A. Wells, Kristin Josic, Maureen G. Maguire, Andrew N. Antoszyk, Carl Baker, Wesley T. Beaulieu, Michael J. Elman, Lee M. Jampol, and Jennifer K. Sun. Five-Year Outcomes after Initial Aflibercept, Bevacizumab, or Ranibizumab Treatment for Diabetic Macular Edema (Protocol T Extension Study). Ophthalmology, 127(9):1201–1210, September 2020.

[33] Alberto Diniz-Filho, Ricardo Y. Abe, Linda M. Zangwill, Carolina P. B. Gracitelli, Robert N. Weinreb, Christopher A. Girkin, Jeffrey M. Liebmann, and Felipe A. Medeiros. Association between Intraocular Pressure and Rates of Retinal Nerve Fiber Layer Loss Measured by Optical Coherence Tomography. Ophthalmology, 123(10):2058–2065, October 2016.

[34] Alka Tripathi, Shagun Gaur, Richa Agarwal, Neha Singh, Amit Singh, Shadman Parveen, Nivedita Singh, and Nani Rima. Disorganization of retinal inner layers as an optical coherence tomography biomarker in diabetic retinopathy: A review. Indian Journal of Ophthalmology, 73(9):1245, September 2025.

