## Supplementary Figures for "Mechanisms of macular oedema development and therapeutic response: An in-silico modelling study"

#### 1 Solid volume fraction map

We observe that, in the physiological case, the solid volume fraction is highest in the macular region (Figure S1a). This is consistent with the physiological understanding that the macula is responsible for the accurate vision, and has a higher density of photoreceptor cells. In the oedematous case, we observe that the solid volume fraction is lower in the macular region compared to the physiological case (Figure S1b). This is consistent with the understanding that in oedema, fluid accumulation leads to swelling and increased interstitial space, reducing the solid volume fraction.

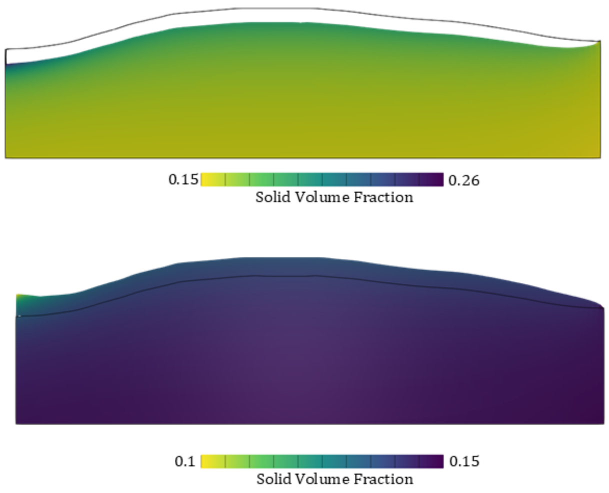


Figure S1: Map of solid volume fraction in the retina, in Physiological (Top) and Oedematous (Bottom) conditions.

#### 2 Diffusion anisotropy vs permeability anisotropy

We studied whether the muller cell orientation affecting diffusion has higher effect on the Anti-VEGF treatment response than the permeability anisotropy. We observed that when both diffusion and permeability are anisotropic, the retinal thickness is highest (Figure S2). When only permeability is anisotropic, the retinal thickness is lower than the previous case. When only Diffusion is anisotropic, we found that the retinal thickness is almost similar to the case when both diffusion and permeability are anisotropic. While the current study shows the non-linear coupling between diffusion and permeability, its clinical significance of needs to be further studied. Global sensitivity analysis can be helpful in understanding the relative importance of these parameters in determining the treatment response.

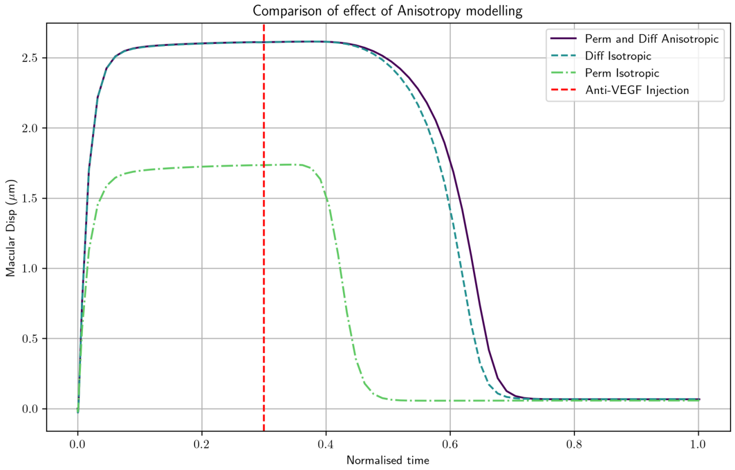


Figure S2: Macular thickness vs time for different combinations of diffusion and permeability anisotropy.

#### 3 IOP vs retinal thicknesses

We studied the effect of intraocular pressure (IOP) on the retinal thickness. We observed that as the IOP increased, more fluid can enter the retina from the vitreous humour, leading to an increase in retinal thickness (Figure S3). This is in contrast with the clinical observation that increased IOP leads to retinal thinning [1]. This discrepancy could be due to the fact that our model does not currently account for the long-term effects of elevated IOP, such as retinal ganglion cell loss and optic nerve damage, which can lead to retinal thinning over time. Future work could involve incorporating these long-term effects into the model to better understand the relationship between IOP and retinal thickness.

#### 4 Protocol T

A longitudinal study on the effect of different anti-VEGF treatment drugs on retinal thickness was performed [2]. Protocol T involves administering anti-VEGF injections and measuring the retinal thickness at various time points. Thy observed that different drugs had different efficacies in reducing retinal thickness over time. Additionally, the reduction in retinal thickness was more pronounced in the initial weeks following treatment,

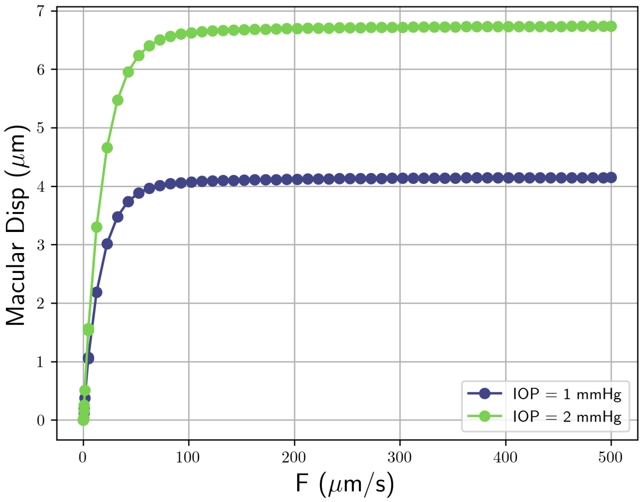


Figure S3: Effect of Intraocular Pressure (IOP) on retinal thickness over time.

37 with a gradual plateauing effect observed over time. This observation was the basis of exponential decay fitting performed in the main manuscript.

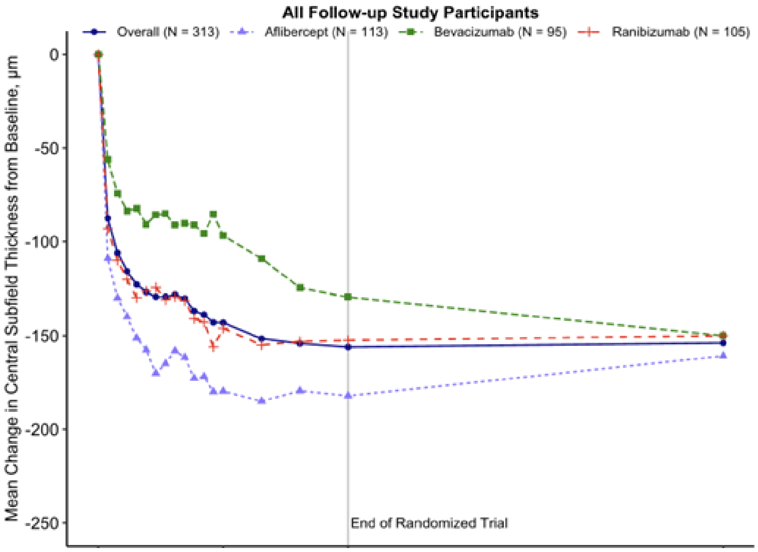


Figure S4: Protocol T anti-VEGF treatment response. Image adapted from [2].

### References

[1] Alberto Diniz-Filho, Ricardo Y. Abe, Linda M. Zangwill, Carolina P. B. Gracitelli, Robert N. Weinreb, Christopher A. Girkin, Jeffrey M. Liebmann, and Felipe A. Medeiros. Association between Intraocular Pressure and Rates of Retinal Nerve Fiber Layer Loss Measured by Optical Coherence Tomography. *Ophthalmology*, 123(10):2058–2065, October 2016.

[2] Adam R. Glassman, John A. Wells, Kristin Josic, Maureen G. Maguire, Andrew N. Antoszyk, Carl Baker, Wesley T. Beaulieu, Michael J. Elman, Lee M. Jampol, and Jennifer K. Sun. Five-Year Outcomes after Initial Aflibercept, Bevacizumab, or Ranibizumab Treatment for Diabetic Macular Edema (Protocol T Extension Study). *Ophthalmology*, 127(9):1201–1210, September 2020.
